# KRAS-driven metabolic reprogramming of enhanced MFSD2A-mediated lysophosphatidylcholine scavenging promotes survival and progression of pancreatic ductal adenocarcinoma

**DOI:** 10.64898/2026.10.05.756824

**Authors:** Yihui Chen, Cason Haffner, Benson Chellakkan Selvanesan, Ranran Wu, Riccardo Ballaro, Ana V. Vilchis Celis, Mingxin Zuo, LaNisha L. Petterson, Michael Adofo, Jody Vykoukal, Ricardo A. León-Letelier, Rongzhang Dou, Soyoung Park, Yining Cai, Daniela B. Rodriquez-Perera, Ehsan Irajizad, Florencia McAllister, Michael Kim, Eugene Koay, Brian Wolpin, Fu Chung Hsiao, Jennifer B. Dennison, Hiroyuki Katayama, Vincent Bernard Pagan, Anirban Maitra, Samir M. Hanash, Johannes F. Fahrmann

## Abstract

Oncogenic KRAS drives lipid reprogramming in pancreatic ductal adenocarcinoma (PDAC) to promote disease progression, and these changes are manifest in circulation. Using mass spectrometry, we performed lipidomic analyses on plasmas from a cohort of 93 newly-diagnosed resectable PDAC cases, 43 patients with chronic pancreatitis, and 93 age- and sex-matched healthy controls, the results of which revealed pronounced reductions in circulating lysophosphatidylcholines (LysoPC) as a prominent feature associated with PDAC. Using stable-isotope resolved lipidomics, we further found that PDAC cells exhibit a lipid scavenging phenotype characterized by preferential uptake of extracellular unsaturated LysoPCs to sustain pro-growth signals. Restricting bioavailability of extracellular unsaturated LysoPCs shifted the metabolic phenotype of PDAC cells towards enhanced *de novo* lipogenesis and autophagy, which was accompanied by suppression of AKT signaling activities and increased susceptibility to apoptosis. Using bulk, single-cell, and spatial transcriptomic datasets together with functional experiments, we identified oncogenic KRAS-mediated upregulation of major facilitator superfamily domain-containing protein 2a (MFSD2A) as a key mediator that facilitates extracellular uptake of LysoPCs in PDAC cells. Loss of MFSD2A expression mimicked LysoPC restriction and resulted in inhibition of AKT signaling and increased propensity of PDAC cells to undergo apoptosis *in vitro* and suppressed tumor growth *in vivo*. Therapeutic targeting of the KRAS-MFSD2A-LysoPC axis via repurposing of the synthetic alkyl-lysophospholipid edelfosine promoted potent anti-cancer effects in vitro that were mechanistically attributed to reduced extracellular uptake of LysoPCs, and attenuation of AKT signaling. Collectively, our findings describe a previously unrecognized onco-metabolic program of enhanced LysoPC scavenging mediated by MFSD2A that promotes PDAC progression and that is therapeutically targetable via repurposing edelfosine.

## Introduction

Pancreatic ductal adenocarcinoma (PDAC) remains among the most lethal solid malignancies with an overall 5-year survival of 13.3% [1, 2]. PDAC is characterized by a high prevalence of activating *KRAS* mutations (∼90–95%) together with recurrent co-alterations in *TP53*, *CDKN2A/B*, and *SMAD4* [3]. These genetic events drive the development of a fibroinflammatory and desmoplastic microenvironment that imposes substantial metabolic stress on cancer cells through hypoxia, fluctuating nutrient availability, and elevated biosynthetic demands [4, 5]. To sustain rapid proliferation under these conditions, KRAS-mutant PDAC cells engage a spectrum of metabolic adaptations that integrate *de novo* synthesis, intracellular recycling pathways, and extracellular nutrient acquisition [6, 7]. In particular, tumor cells become increasingly reliant on exogenous lipid sources to sustain membrane synthesis, signaling, and survival under metabolic stress [7, 8]. Consistent with this concept, disruption of extracellular lipid supply or intracellular lipid processing exposes compensatory metabolic programs that can be therapeutically targeted in preclinical PDAC models [9, 10]. Importantly, the abundance and composition of extracellular lipids are determined not only by local tumor and stromal metabolism, but also by systemic lipid supply [8, 11]. In mouse models of PDAC, dietary manipulation alters lipid abundance in both plasma and the tumor microenvironment and influences tumor progression, highlighting a functional link between host lipid metabolism and tumor lipid utilization [11]. These observations suggest that changes in circulating lipids may not simply represent biomarkers associated with presence of disease, but may also reflect metabolic dependencies that support tumor growth. Defining such alterations in the circulating lipidome could therefore provide insight into nutrient acquisition strategies used by PDAC cells and reveal actionable metabolic vulnerabilities.

In the current study, we performed lipidomic analyses on plasma samples from a cohort of 93 patients with early-stage PDAC, 43 patients with chronic pancreatitis (CP), and 93 age- and sex-matched healthy controls to characterize PDAC-associated changes in circulating lipid profiles. These analyses identified reduced circulating LysoPC levels as a prominent metabolic feature associated with PDAC. Using an integrated approach, we found that oncogenic KRAS upregulates cell-surface expression of major facilitator superfamily domain-containing protein 2a (MFSD2A), promoting preferential uptake of unsaturated LysoPC species that sustain AKT signaling and suppress apoptosis. Moreover, we demonstrate that this KRAS-MFSD2A-LysoPC axis represents a metabolic vulnerability that can be therapeutically targeted via repurposing of the synthetic LysoPC analog edelfosine.

## Results

### Reductions in circulating lysophosphatidylcholine (LysoPC) species are a prominent feature associated with PDAC

To interrogate alterations in circulating lipid profiles associated with PDAC, we performed lipidomic analyses of plasma samples from a retrospective cohort of 99 patients with early-stage PDAC, 46 patients with chronic pancreatitis (CP), and 96 age- and sex-matched healthy controls. A total of 247 annotated lipids were identified, for which 75 were found to differ significantly (Wilcoxon rank sum test 2-sided p-value <0.05) between cases compared to respective controls (healthy plus patients with CP). Among the altered lipid classes, lysophospholipids, particularly lysophosphatidylcholine (LysoPCs), emerged as one of the most consistently depleted lipid groups in patients with PDAC relative to both healthy controls and CP patients (**Figure 1A-C, Supplementary Figure S1A-1B, and Supplementary Table S1**). Multiple LysoPC species, especially the long-chain unsaturated long-chain and very-long-chain LysoPCs, exhibited significantly lower abundance in PDAC plasma (Welch’s t test 2-sided p-value <0.05, **Figure 1D**). Prior studies by our group and others have similarly documented reductions in circulating LysoPCs as a metabolic manifestation associated with presence of PDAC [12, 13, 14].

**Figure 1.**
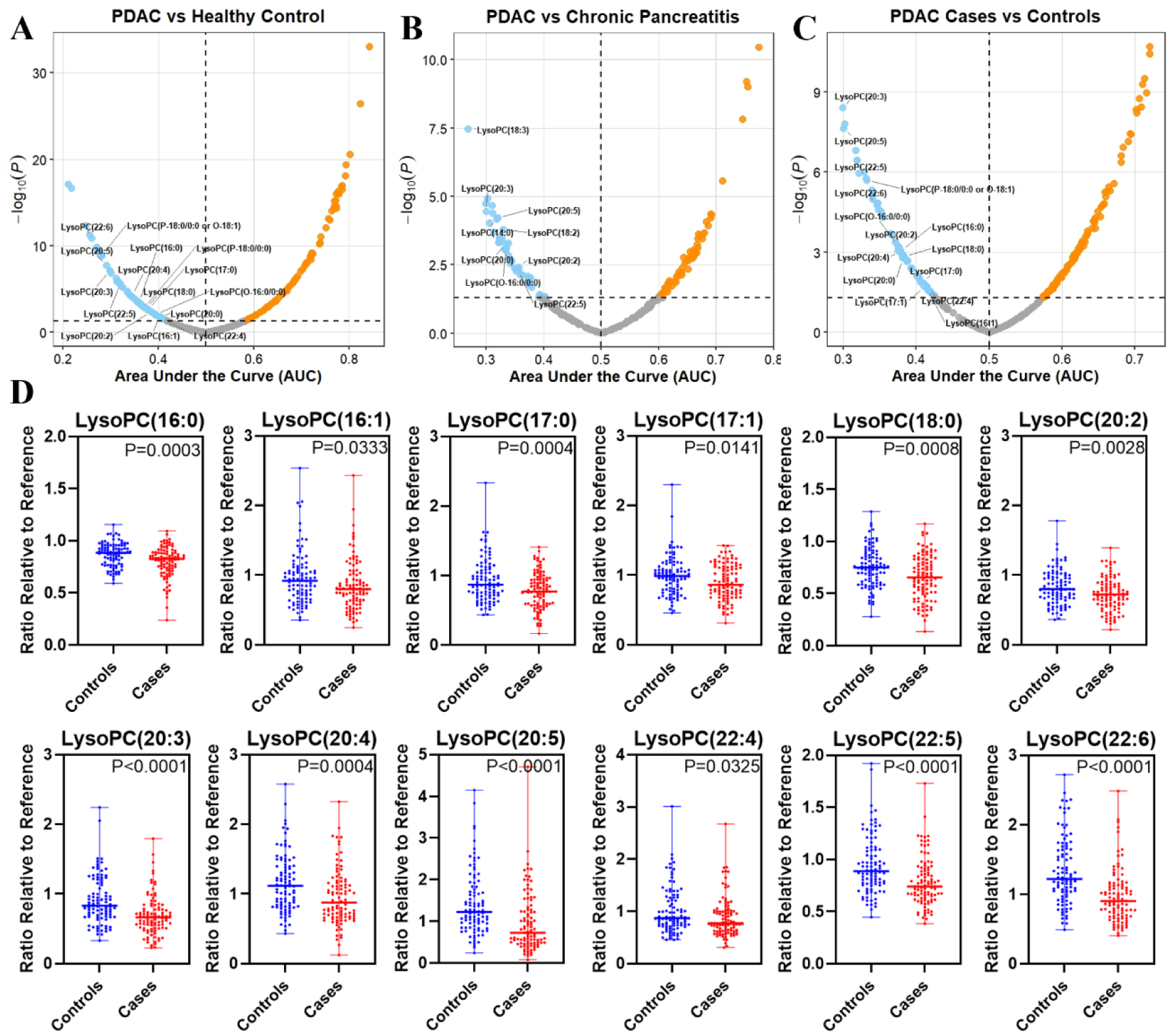
Circulating LysoPCs are reduced in PDAC. Lipidomic profiling of plasma samples from patients with newly-diagnosed resectable PDAC (n=99), healthy controls (n=96), and patients with chronic pancreatitis (CP, n=46). Differential lipid analyses were performed by comparing PDAC cases with healthy controls **(A)**, PDAC cases with CP patients **(B)**, and PDAC cases with all controls combined (healthy controls plus CP patients; **C).** Volcano plots display the discriminatory performance of individual lipid species, where each point represents a lipid. Statistical significance was determined by two-sided Wilcoxon rank-sum tests. **D)** Distribution plots illustrating relative abundance (compared to reference historical quality control) of LysoPC species in plasmas of PDAC cases (n=99) and healthy controls (n=96). Statistical significance was determined by Welch’s t test and 2-sided p values are reported.

### Pancreatic cancer cells scavenge extracellular LysoPCs to sustain proliferation under nutrient deprivation

We considered that the reduction in circulating lysophospholipids may reflect enhanced scavenging and catabolism by PDAC cells. To test this hypothesis, we first performed lipidomic analyses on conditioned medium (CM) temporally collected from cultured human PDAC and murine KPC cell lines and confirmed rapid depletion of LysoPCs and other lysophospholipid species (**Figure 2A**). In contrast, other lipid classes, e.g. phosphatidylcholines (PCs) and triacylglycerol (TAGs), demonstrated variable depletion across the evaluated human and murine PDAC cell lines (**Supplementary Table S2**).

**Figure 2.**
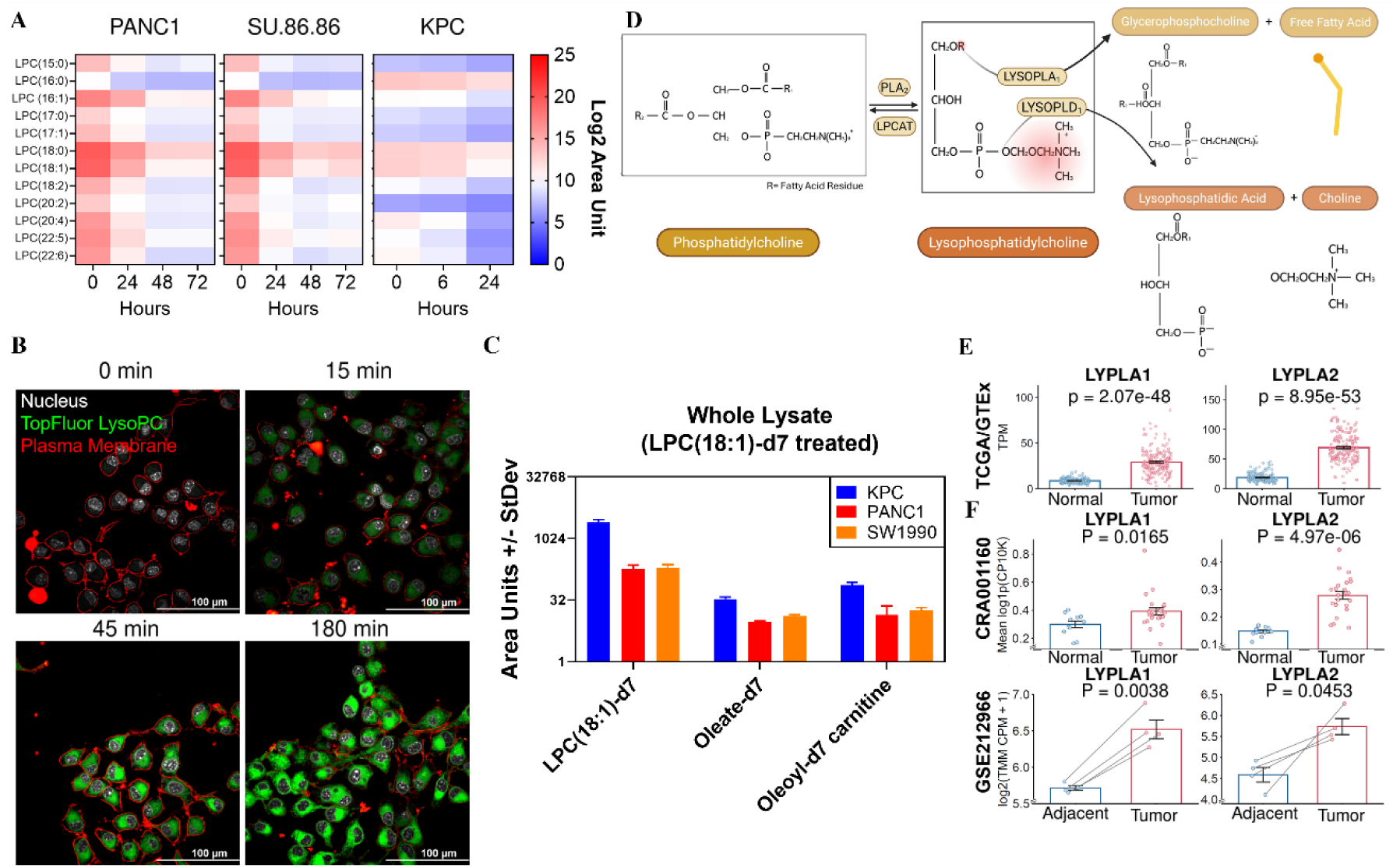
Pancreatic cancer cells readily scavenge extracellular LysoPCs. **A)** Heatmap illustrating abundance of LysoPC species in conditioned medium collected from human PDAC and murine KPC cell lines. **B)** Confocal of LysoPC uptake (TopFluor LysoPC, 1 µM). **C)** Abundance of isotope labeled LysoPC species and related lipids in human PDAC and murine KPC cells treated with deuterated LysoPC(18:1)-d7 (1 µg/ml). **D)** Schematic of LysoPC metabolic fate. **E)** Expression levels of *LYPLA1* and *LYPLA2* in normal pancreas (GTEx) and PDAC (TCGA) tissues. Statistical significance was determined by Wilcoxon rank-sum test and 2-sided p values are reported. **F)** Expression levels of *LYPLA1* and *LYPLA2* in epithelial/ductal cells from normal pancreas and PDAC tissues. Statistical significance was determined by Wilcoxon rank-sum test (CRA001160) and paired t-test (GSE212966), with 2-sided p values reported.

We next sought to determine whether the observed reductions in CM lysophospholipids were attributed to enhanced uptake by PDAC cells or catabolism by secreted lipases. For these studies we focused on LysoPCs as these are the most abundant type of lysophospholipids in circulation. Treatment of KPC cells with fluorescently labeled TopFluor-LysoPC followed by confocal microscopy confirmed rapid internalization (**Figure 2B**). Similarly, treatment of PDAC cell lines (PANC1, SW1990, KPC) with deuterated LysoPC(18:1)-d7 followed by mass spectrometry-based analysis of cell extracts revealed detectable levels of LysoPC(18:1)-d7 (**Figure 2C**), reinforcing that PDAC cells consume extracellular LysoPCs.

LysoPC can be metabolized through multiple pathways, including metabolism to lysophosphatidic acid (LysoPA) and choline by lysophospholipase D or hydrolysis into glycerophosphocholine (GPC) and free fatty acids (FFA) by lysophospholipase A (**Figure 2D**). Using stable-isotope data generated on LysoPC(18:1)-d7 treated PDAC cells, we screened for downstream isotopically-labeled LysoPC metabolites. These analyses revealed detectable levels of oleate-d7 and oleoyl-d7-carnitine (**Figure 2C**). We did not observe detectable levels of LPA(18:1)-d7. Collectively, these data indicate hydrolysis of intracellular LysoPCs by lysophospholipase A and shunting of liberated FFA towards downstream fatty-acid metabolism. In support of these findings, mRNA expression levels of key lysophospholipase A enzymes, including LYPLA1 and LYPLA2, in PDAC tumors and normal pancreas tissues from the TCGA and GTEx revealed substantial upregulation in PDAC tumors (**Figure 2E and Supplementary Figure S2A**). Similar findings were found in epithelial/ductal cell populations from independent single-cell transcriptomic datasets (CRA001160 and GSE212966, **Figure 2F and Supplementary Figure S2B and S2C**).

Environmental lipids represent an important nutrient source for PDAC cells [6]. To determine the consequences of lipid restriction broadly, we cultured human PDAC (PANC1 and SU.86.86) and murine KPC cells in delipidated medium. The results of these studies revealed pronounced growth inhibition and increased apoptosis, as measured by caspase-3/7 activity, compared to lipid-containing medium (**Supplementary Figure S3A and S3B**). Lipid deprivation also induced compensatory autophagy, as demonstrated by increased p62-LC3B accumulation and colocalization, and upregulation of fatty acid synthase (FASN) and sterol regulatory element-binding protein 1 (SREBP1) protein expression, indicating activation of the *de novo* lipogenesis pathway (**Supplementary Figure S3C and S3D)**. Moreover, extracellular lipid restriction sensitized PDAC cells to pharmacologic inhibitors of autophagy (Azithromycin, Bafilomycin A1, and Chloroquine) and fatty acid synthesis (Denifanstat) (**Supplementary Figure S3E and S3F**).

To explicitly determine dependency of PDAC cells on extracellular LysoPCs, we supplemented human PANC1, SU.86.86, and murine KPC cells cultured in delipidated medium with LysoPC(18:0), LysoPC(18:1), or LysoPC(18:2). As comparative control, we also included phosphatidylcholine(18:1,18:1) (hereon referred to as DOPC). Supplementation with unsaturated LysoPC species (LysoPC(18:1) and (18:2)) restored cell proliferation across both human and murine PDAC cell lines (**Figure 3A**). Similar growth-promoting effects were observed following supplementation with other unsaturated lysophospholipids (**Supplementary Figure S4A**). In contrast, LysoPC(18:0) exhibited dose-dependent toxicity consistent with the known lipotoxic effects of saturated lysophospholipids [8, 15, 16] (**Figure 3A**). DOPC produced variable and cell line-dependent effects. Consistent with their effects on proliferation, unsaturated LysoPCs, as shown with LysoPC(18:1), reduced apoptosis, decreased reliance of PDAC cells on autophagy and *de novo* lipogenesis, and reduced sensitivity to autophagy and FASN inhibitors (**Figure 3B-F**).

**Figure 3.**
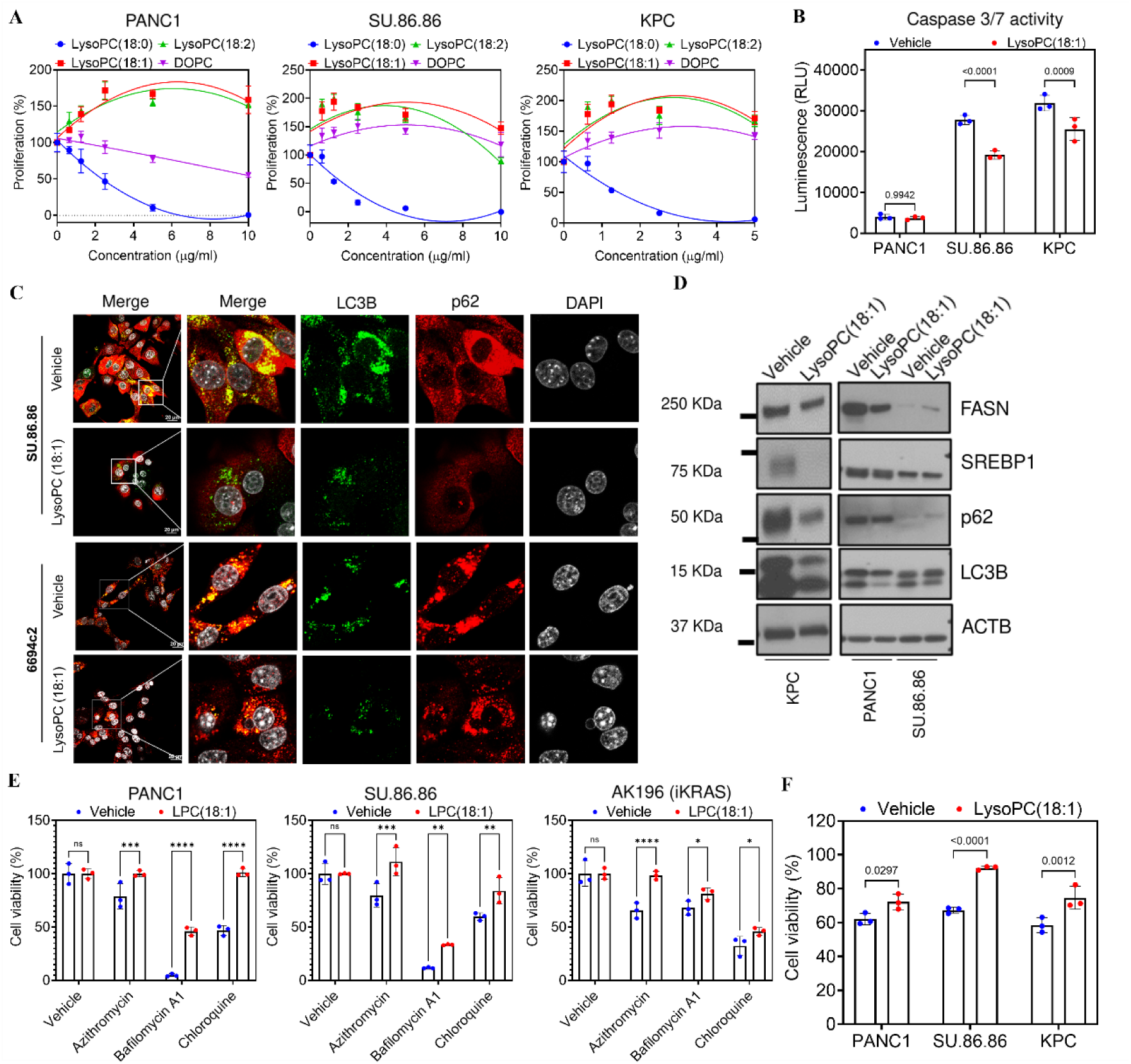
Salvaging Extracellular Unsaturated LysoPCs Maintains Lipid Homeostasis and Promotes PDAC Cell Proliferation and Survival. **A)** Cell viability of human PDAC and murine KPC cells treated with LysoPC(18:0), LysoPC(18:1), LysoPC(18:2), or DOPC at the concentrations indicated. **B)** Caspase 3/7 activity in human PDAC and murine KPC cells treated with vehicle control or LysoPC(18:1) (10 µg/ml). Statistical significance was determined by Šídák’s multiple comparisons test. **C)** Confocal imaging of LC3B (Green), p62 (Red) and nucleus (white) in human PDAC and murine KPC cells treated with vehicle control or LysoPC(18:1) (10 µg/ml). **D)** Immunoblotting of FASN, SREBP1, p62, LC3B, and ACTB in human PDAC and murine KPC cells treated with vehicle control or LysoPC(18:1) (10 µg/ml). **E)** Cell viability of PANC1, SU.86.86, and AK196 (iKRAS) cells treated with vehicle control or LysoPC(18:1) (10 µg/ml) with autophagy inhibitors. Statistical significance was determined by Fisher’s LSD test. **F)** Cell viability of human PDAC and murine KPC cells treated with vehicle control or LysoPC(18:1) (10 µg/ml) with FASN inhibitor. Statistical significance was determined by Šídák’s multiple comparisons test.

Previous studies have shown that LysoPC can be converted into biologically active metabolites, including LysoPA and free fatty acids (**Figure 2D**), which themselves can influence tumor growth [17]. To determine whether metabolic byproducts derived from metabolism of unsaturated LysoPC would similarly elicit pro-growth effects, we treated human (PANC1 and SU.86.86) and murine (KPC) PDAC cells with LysoPC(18:1), LysoPA(18:1) plus choline, or GPC plus FFA(18:1). Although all experimental conditions promoted cancer cell growth and blunted apoptosis, LysoPC(18:1) consistently elicited the strongest effects (**Figure 4A and 4B**). These findings insinuate that intact LysoPC possesses biological activity beyond simply serving as a precursor for downstream metabolites. These observations are also consistent with prior reports demonstrating that direct blocking of autotaxin-LysoPA axis or LysoPA receptors only minimally reduced LysoPC-mediated cell growth [8].

**Figure 4.**
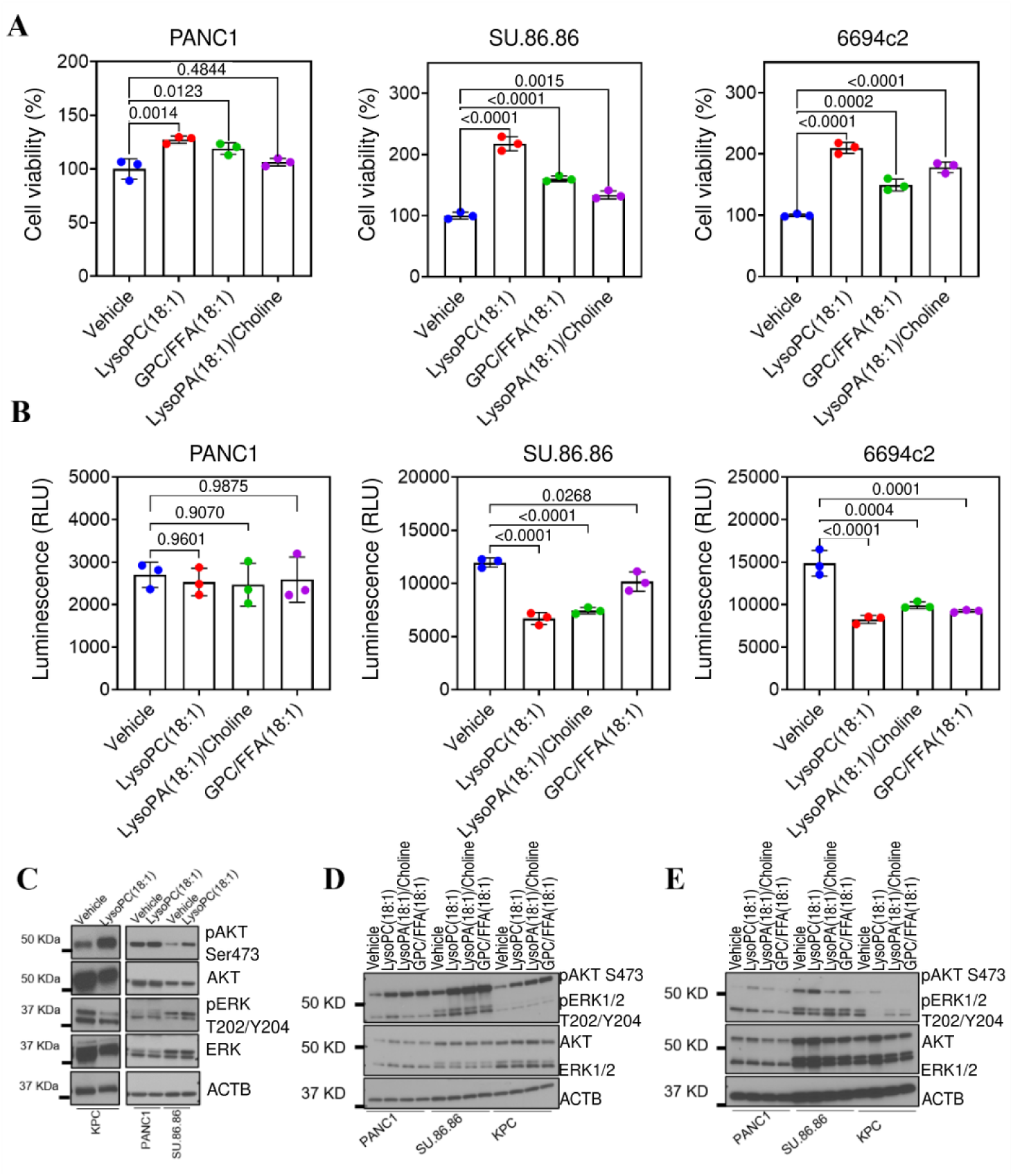
LysoPC supports PDAC cancer cell growth and oncogenic signaling independent from its metabolic products. **A)** Cell viability of human PDAC and murine KPC cells treated with vehicle control, LysoPC(18:1), LysoPA(18:1)/Choline, or GPC/FFA(18:1) (5 µM). Statistical significance was determined by Dunnett’s multiple comparisons test. **B)** Caspase 3/7 activity in human PDAC and murine KPC cells treated with vehicle control, LysoPC(18:1), LysoPA(18:1)/Choline, or GPC/FFA(18:1) (5 µM). Statistical significance was determined by Tukey’s multiple comparisons test. **C)** Immunoblotting of phosphorylated AKT (pAKT, S473), AKT, pERK1/2 (T202/Y204), ERK1/2, and ACTB in human PDAC and murine KPC cells treated with vehicle control or LysoPC(18:1) (10 µM). The ACTB panel is the same as Figure 3D. **D)** Immunoblotting of phosphorylated AKT (pAKT, S473), AKT, pERK1/2 (T202/Y204), ERK1/2, and ACTB in human PDAC and murine KPC cells treated with vehicle control, LysoPC(18:1), LysoPA(18:1)/Choline, or GPC/FFA(18:1) (10 µM) for 15 minutes. **E)** Immunoblotting of phosphorylated AKT (pAKT, S473), AKT, pERK1/2 (T202/Y204), ERK1/2, and ACTB in human PDAC and murine KPC cells treated with vehicle control, LysoPC(18:1), LysoPA(18:1)/Choline, or GPC/FFA(18:1) (10 µM) for 48 hours.

### Unsaturated LysoPCs sustain PI3K-AKT signaling to promote growth of PDAC cells

We considered that the observed pro-growth effects of unsaturated LysoPCs may be attributed to PI3K-AKT signaling activities, which exhibit bi-directional relationships with lipid metabolism [17]. Consistent with this hypothesis, supplementation of PANC1, SU.86.86 and KPC cells cultured in delipidated media with LysoPC(18:1) or LysoPC(18:2), but not LysoPC(18:0) or DOPC, enhanced phosphorylation of AKT at Ser473 (**Figure 4C, Supplementary Figure S4B and S4C**). Acute treatment (15 minutes) of cells with LysoPC(18:1), GPC/FFA(18:1), or LysoPA(18:1)/choline led to comparable increase in AKT phosphorylation (**Figure 4D**). However, following prolonged exposure (48 hours), sustained AKT activation was observed only in cells treated with intact LysoPC(18:1) (**Figure 4E**). These findings suggest that while LysoPC metabolites can contribute to acute signaling responses, intact LysoPC supports durable pro-survival AKT signaling.

Collectively, these results demonstrate that extracellular unsaturated LysoPCs function as metabolically important lipid nutrients in PDAC. By supporting lipid homeostasis, suppressing adaptive stress responses, and sustaining AKT signaling, LysoPC scavenging program promotes tumor cell proliferation and survival.

### Oncogenic KRAS-mediated upregulation of MFSD2A promotes LysoPC uptake in pancreatic cancer cells

Activating KRAS mutations occur in approximately 90% of PDACs and are key drivers of tumor initiation, progression, and metabolic adaptation [3, 8, 9, 10, 18]. Prior study has also suggested that oncogenic signaling, including RAS activation, drives the selective salvaging of extracellular unsaturated fatty acids derived from catabolism of lysophospholipids [7]. Recently, major facilitator superfamily domain-containing protein 2A (MFSD2A), also known as sodium-dependent lysophosphatidylcholine symporter 1, was identified as a key transporter that mediates uptake of extracellular LysoPCs in other diseases [19, 20].

Analysis of transcriptomic data from TCGA and GTEx similarly revealed statistically significantly higher *MFSD2A* expression in PDAC tissues relative to normal pancreas (**Figure 5A**). Similar results were observed in an independent dataset containing 118 PDAC and 13 normal pancreatic tissues (GSE62165; **Figure 5B**) [21]. Moreover, *MFSD2A* expression was statistically significantly higher in KRAS-mutant tumors than in KRAS wild-type PDACs (TCGA, **Figure 5C**). Evaluation of independent single-cell RNA-sequencing datasets (CRA001160 and GSE212966) further demonstrated enrichment of *MFSD2A* within malignant epithelial/ductal cell populations (**Figure 5D**) [22, 23]. Consistent with these observations, spatial transcriptomic analyses localized *MFSD2A* expression predominantly to tumor epithelial regions (GSE272362, **Figure 5E-F**) [24]. Notably, elevated *MFSD2A* expression was also observed in hypoxic tumor regions (**Supplementary Figure S5**), consistent with a potential role for MFSD2A-mediated lipid acquisition under metabolically stressful conditions.

**Figure 5.**
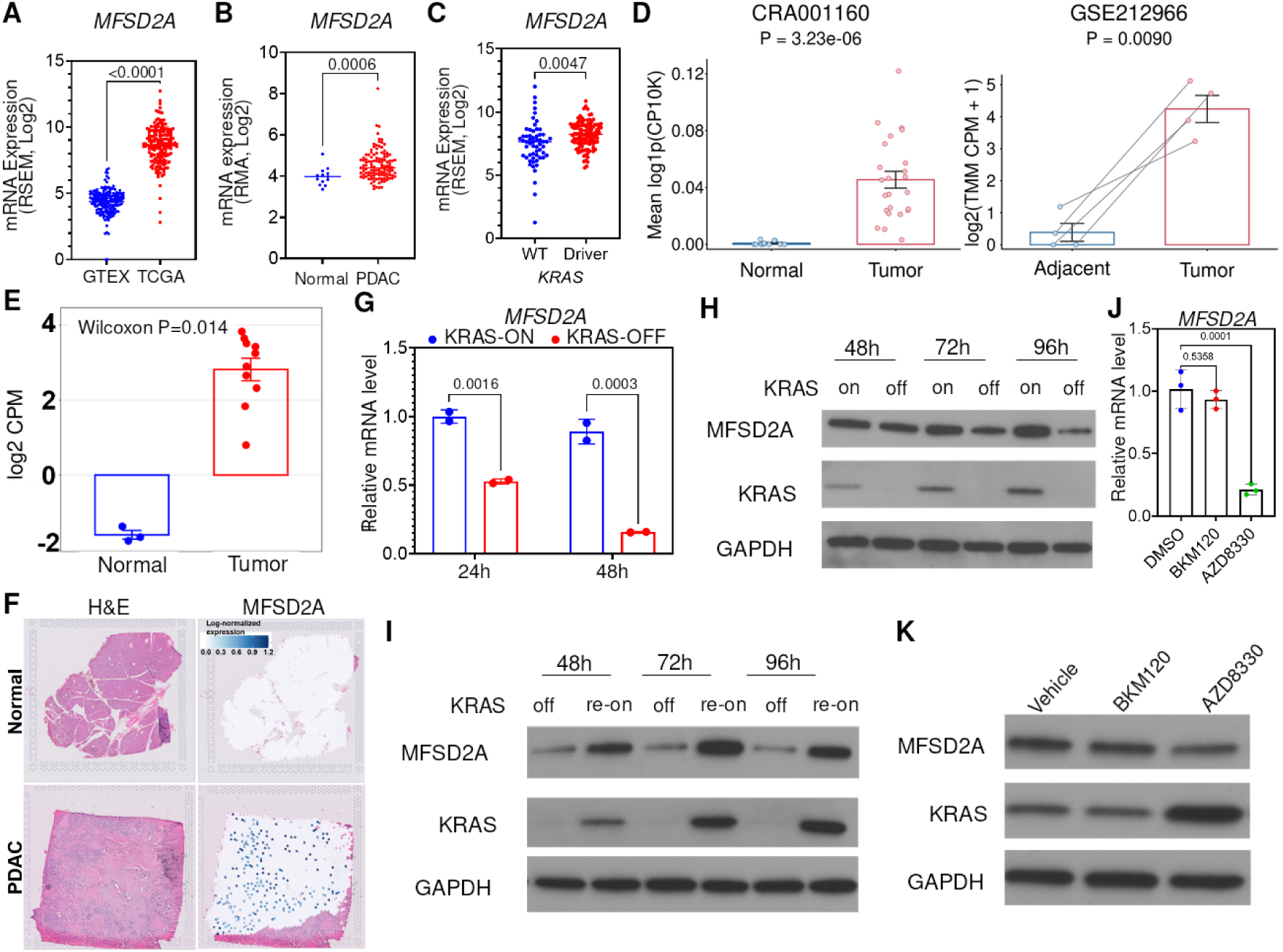
Oncogenic KRAS signaling regulates *MFSD2A* expression. **A-B**) Expression levels of MFSD2A in GTEx and TCGA PAAD tissues (**A**) and in normal pancreatic tissues and PDAC tumor tissues from GSE62165 (**B**). Statistical significance was determined by unpaired t-test. **C)** mRNA expression level of MFSD2A in PDAC tumor tissues with KRAS WT or driver mutations from TCGA-PAAD dataset. Statistical significance was determined by unpaired t-test. **D)** Expression levels of MFSD2A in epithelial/ductal cells in adjacent normal pancreas and PDAC tissues. Data was obtained from CRA001160 (left panel) and GSE212966 (right panel). Statistical significance was determined by Wilcoxon rank-sum test (CRA001160) and paired t-test (GSE212966). **E)** Expression levels of MFSD2A in normal pancreas and PDAC tissues. Data was obtained from GSE272362. Statistical significance was determined by Wilcoxon rank-sum test and 2-sided p values are reported. **F)** Spatial distribution of MFSD2A in normal and PDAC tissues. Data was obtained from GSE272362. **G)** mRNA expression level of MFSD2A in iKRAS cells with or without DOX (1 µg/ml). Statistical significance was determined by Šídák’s multiple comparisons test. **H)** Immunoblots for MFSD2A and KRAS in iKRAS cells after withdrawal of DOX (1 µg/ml) for the indicated times. **I)** Immunoblots for MFSD2A and KRAS in iKRAS cells cultured in DOX-free medium for 24 hours and re-challenged with DOX (1 µg/ml) for indicated time. **J-K)** mRNA (**J**) and protein (**K**) levels of MFSD2A in KPC cells treated with DMSO, PI3K inhibitor (BKM120) or MEK inhibitor (AZD8330). Statistical significance was determined by Dunnett’s multiple comparisons test.

To determine whether KRAS signaling directly regulates MFSD2A, we utilized inducible KRAS (iKRAS) PDAC cells, in which oncogenic KRAS activity can be reversibly controlled [25]. Suppression of KRAS signaling resulted in a progressive reduction of MFSD2A mRNA and protein expression (**Figure 5G-H**), whereas reactivation of KRAS restored MFSD2A expression (**Figure 5I**). Pharmacological interrogation of KRAS effector pathways revealed that inhibition of MAPK signaling markedly reduced MFSD2A expression, whereas blockade of alternative downstream pathways had comparatively limited effects (**Figure 5J-K**).

To elucidate the reliance of PDAC cells on MFSD2A-mediated uptake of LysoPCs, we generated *MFSD2A*-deficient SU.86.86 and KPC cells using shRNA-mediated knockdown (**Figure 6A**). Lipidomic analyses revealed markedly delayed depletion of both LysoPC(18:1)-d7 and LysoPC species in CM of shMFSD2A PDAC cells compared to shCTRL cells (**Figure 6B-C**). In contrast, abundances of other lipid classes remained largely unchanged (**Supplementary Table S3**). Importantly, MFSD2A knockdown disproportionately attenuated the depletion of unsaturated LysoPCs (**Figure 6C**), indicating preferential selectivity of MFSD2A towards unsaturated LysoPCs [19]. Independent assessment of LysoPC uptake using fluorescently labeled TopFluor-LysoPC similarly demonstrated reduced intracellular accumulation in shMFSD2A PDAC cells compared to shControl (**Figure 6D-E**). Moreover, isotope-tracing studies revealed reduced generation of FA(18:1)-d7 from LysoPC(18:1)-d7 in MFSD2A-deficient cells (**Figure 6F**), indicating impaired intracellular processing of imported LysoPC.

**Figure 6.**
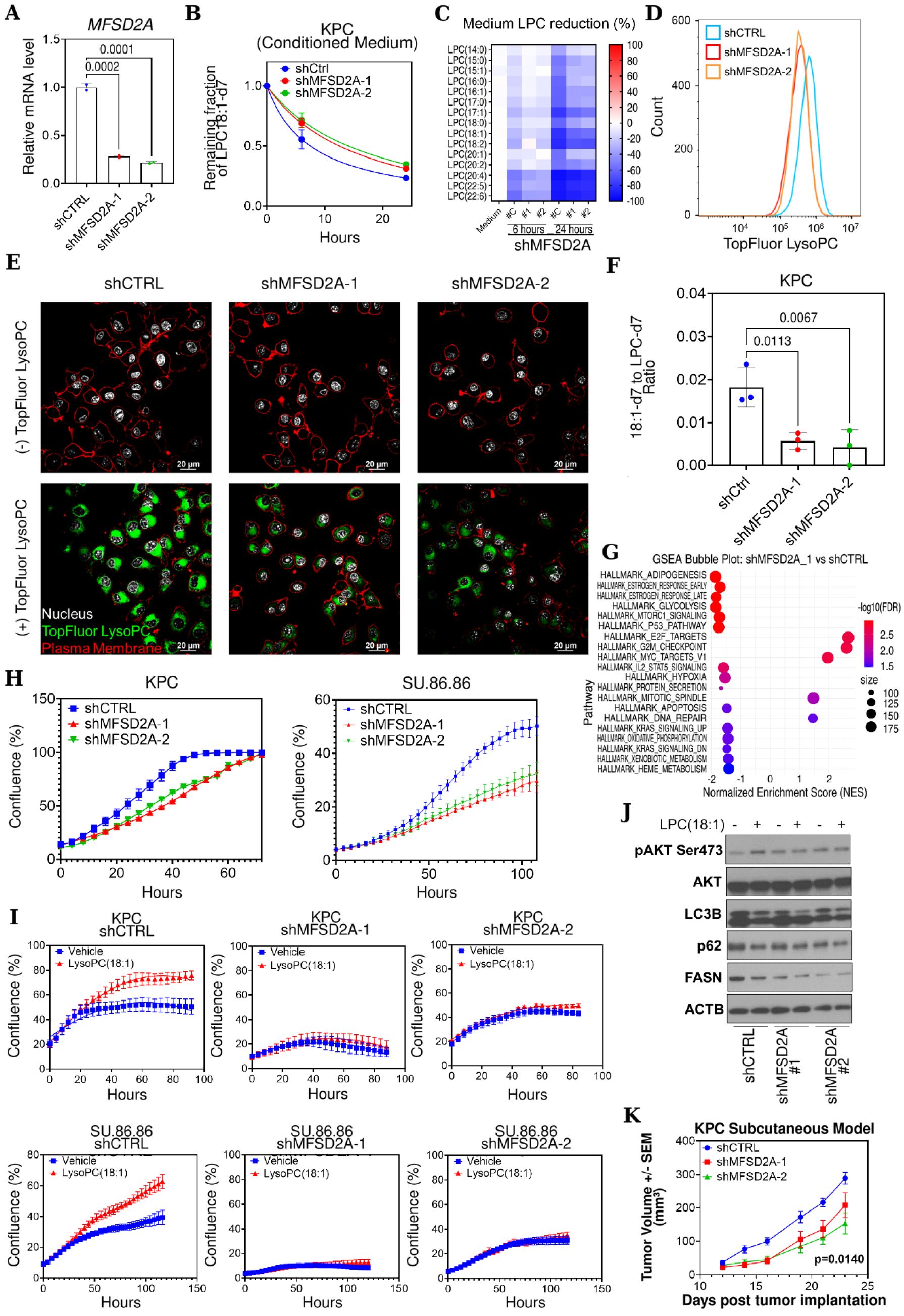
MFSD2A mediates LysoPC uptake in pancreatic cancer cells. **A)** qPCR quantification of mRNA levels of *MFSD2A* in shCTRL and shMFSD2A KPC cells. Statistical significance was determined by Dunnett’s multiple comparisons test. **B)** Abundance of LysoPC(18:1)-d7 in conditioned medium collected from murine shCTRL and shMFSD2A KPC cells. **C)** Percentage reduction of LysoPC species in conditioned medium collected from murine shCTRL and shMFSD2A KPC cells. **D)** Flow cytometry analysis of LysoPC uptake after 3-hour treatment with TopFluor LysoPC (1 µM). **E)** Confocal of LysoPC uptake after 3-hour treatment with TopFluor LysoPC (1 µM). **F**) Relative levels of FFA(18:1)-d7 to LysoPC(18:1)-d7 in murine shCTRL and shMFSD2A KPC cells. Statistical significance was determined by Dunnett’s multiple comparisons test. **G**) GSEA analysis of hallmark pathways in murine shCTRL and shMFSD2A KPC cells. **H)** Impedance-based assay of cell confluence of human and murine shCTRL and shMFSD2A cells. **I)** Impedance-based assay of cell confluence of human and murine shCTRL and shMFSD2A cells treated with vehicle controls or LysoPC(18:1) (10 µg/ml). **J)** Immunoblotting of phosphorylated AKT (pAKT, S473), AKT, LC3B, p62, FASN, and ACTB in murine shCTRL and shMFSD2A KPC cells treated with vehicle control or LysoPC(18:1) (10 µg/ml). **K)** Tumor volume (Mean ± standard error of mean (SEM)) of subcutaneous model of murine shCTRL and shMFSD2A KPC cell. Statistical significance was evaluated using two-way repeated-measures ANOVA with Geisser-Greenhouse correction, followed by Dunnett’s multiple-comparisons test against shCTRL.

We next performed bulk RNA-sequencing analyses of shCTRL and shMFSD2A KPC cells to interrogate signaling activities mediated by the KRAS-MFSD2A-LysoPC axis. Gene set enrichment analysis (GSEA) revealed elevation of cell cycle arrest-associated programs together with reduced PI3K-AKT signaling following MFSD2A knockdown (**Figure 6G and Supplementary Figure S6**). Consistent with these transcriptional changes, MFSD2A-deficient cells exhibited significantly reduced proliferation compared with control cells (**Figure 6H**). Importantly, supplementation with LysoPC(18:1) in delipidated medium stimulated proliferation of shCTRL cells but failed to rescue growth in shMFSD2A cells (**Figure 6I**), indicating that MFSD2A is required for the growth-promoting effects of extracellular LysoPCs. Likewise, LysoPC(18:1)-induced AKT phosphorylation (Ser473) was drastically reduced in MFSD2A KD compared to shControl KPC cells (**Figure 6J**). *In vivo*, MFSD2A-deficient tumors exhibited significantly reduced growth relative to shControl tumors (two-way repeated-measures ANOVA, p=0.0140, **Figure 6K**). Together, these data demonstrate that MFSD2A-dependent LysoPC uptake supports AKT signaling, tumor cell proliferation, and PDAC growth. Collectively, these data identify MFSD2A as a KRAS-regulated lipid transporter and establish a mechanistic link between oncogenic KRAS signaling and enhanced LysoPC scavenging in PDAC cells.

### The KRAS-MFSD2A-LysoPC axis is an actionable metabolic vulnerability in PDAC

Our findings indicate that the KRAS-MFSD2A-LysoPC axis plays a critical role in supporting PDAC growth and survival, raising the possibility that this pathway may be therapeutically targetable. Alkyl-lysophospholipids (ALPs) are synthetic LysoPC analogs in which the ester-linked fatty acyl chain is replaced by an ether-linked alkyl chain, rendering them resistant to enzymatic degradation and metabolically stable. Several ALPs, most notably edelfosine, have demonstrated potent antitumor activity in preclinical models of both hematologic malignancies and solid tumors [26, 27]. However, clinical translation of this drug class has produced mixed results [28, 29, 30], suggesting that their efficacy may depend on the presence of specific metabolic determinants.

Given our findings that MFSD2A mediates LysoPC uptake, we hypothesized that MFSD2A may also be a key determinant of anti-cancer efficacy of edelfosine. To this end, analysis of CM revealed that reduction of edelfosine was significantly attenuated in shMFSD2A cells compared to shCTRL PDAC cells (**Figure 7A)** [19, 31].

**Figure 7.**
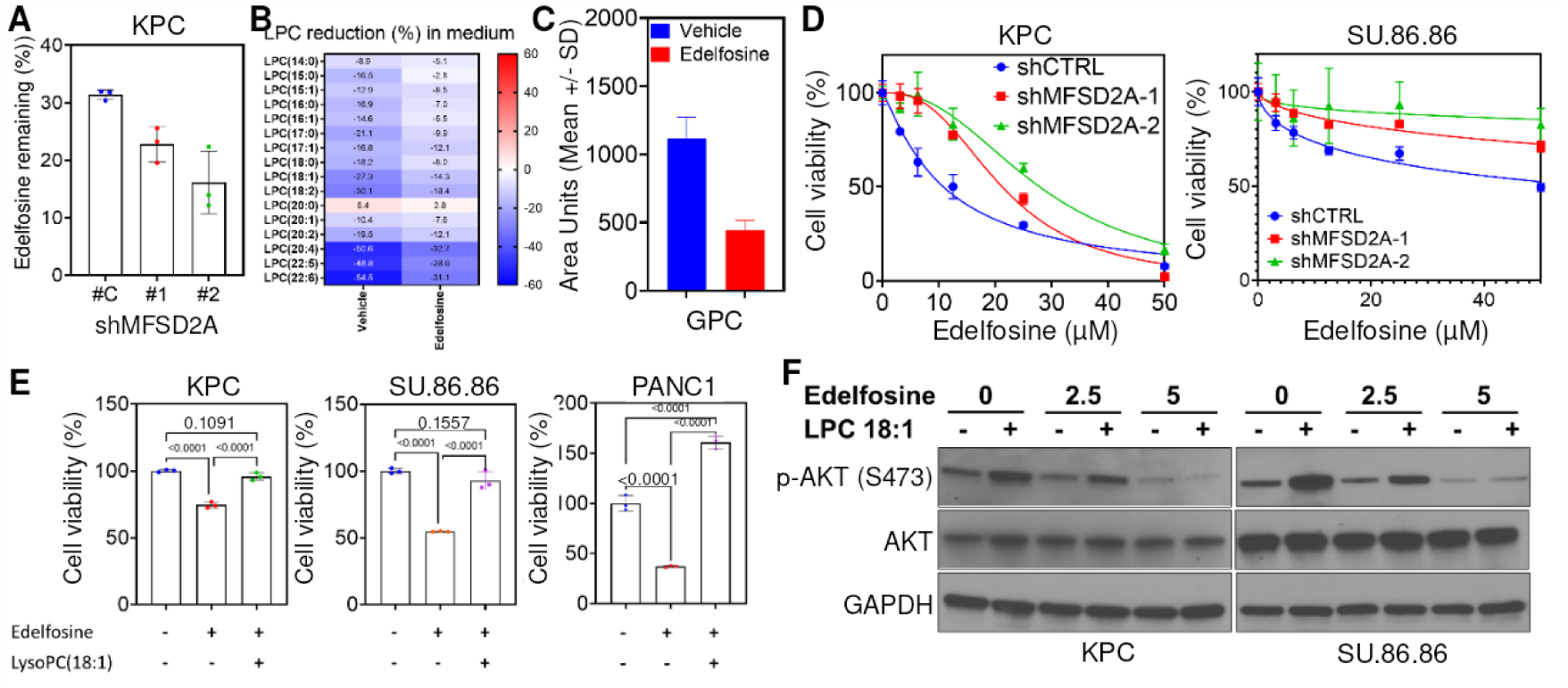
The KRAS-MFSD2A-LysoPC axis is an actionable metabolic vulnerability in PDAC. **A)** Abundance of edelfosine in conditioned medium collected from murine shCTRL and shMFSD2A KPC cells. Statistical significance was determined by Dunnett’s multiple comparisons test. **B)** Percentage reduction of LysoPC species in conditioned medium collected from murine KPC cells treated with vehicle control or edelfosine. **C)** Abundance of GPC in conditioned medium collected from murine KPC cells treated with vehicle control or edelfosine. **D)** Cell viability of KPC and SU.86.86 shCTRL and shMFSD2A cells treated with edelfosine. **E)** Cell viability of human PDAC and murine KPC cells treated with vehicle control, edelfosine, or edelfosine + LysoPC(18:1). Statistical significance was determined by Tukey’s multiple comparisons test. **F)** Immunoblots for p-AKT (S473), AKT, and GAPDH in KPC and SU.86.86 cells treated vehicle control, or LysoPC(18:1) and edelfosine.

Given the structural similarity between edelfosine and LysoPC, we next examined whether edelfosine interferes with LysoPC uptake and utilization by PDAC cells. To this end, treatment of KPC cell lines with edelfosine reduced the depletion of extracellular LysoPC species from CM and suppressed downstream LysoPC metabolism, including conversion into GPC (**Figure 7B-C**). Consistent with MFSD2A-dependent uptake, MFSD2A-deficient cells exhibited reduced sensitivity to edelfosine compared with control cells (**Figure 7D**). These findings indicate that edelfosine disrupts cellular utilization of extracellular LysoPCs.

To further evaluate the functional relationship between edelfosine and LysoPC, PANC1, SU.86.86 and KPC cells were co-supplemented with LysoPC(18:1) during edelfosine treatment. Exogenous LysoPC(18:1) partially rescued cell viability in the presence of edelfosine (**Figure 7E**), indicating that the two molecules functionally antagonize one another. Mechanistically, edelfosine abrogated LysoPC(18:1)-induced AKT phosphorylation (Ser473) in a dose-dependent manner (**Figure 7F**), demonstrating that edelfosine suppresses a key downstream signaling pathway activated by extracellular LysoPCs.

Collectively, these findings identify MFSD2A-mediated LysoPC uptake as a targetable metabolic process in PDAC and provide evidence that the synthetic LysoPC analog edelfosine can disrupt this pathway. These results further suggest that tumors with elevated KRAS-MFSD2A-LysoPC signaling may be particularly susceptible to therapeutic strategies that interfere with LysoPC acquisition and utilization.

## Discussion

Oncogenic KRAS signaling orchestrates massive metabolic reprograming in PDAC to sustain growth within a nutrient-deprived and hypoxic tumor microenvironment [4, 5]. Among these adaptations, lipid metabolism is maintained through a coordinated network of *de novo* synthesis, intracellular recycling, and extracellular acquisition [9, 10, 18]. Because oxygen availability, reducing equivalents, and lipogenic carbon substrates are frequently limited in PDAC, tumor cells increasingly rely on exogenous lipids to support membrane biogenesis, energy metabolism, and pro-survival signaling [6, 7, 8, 32, 33, 34, 35]. In the present study, we identify a previously unrecognized KRAS-MFSD2A-LysoPC axis that links oncogenic signaling to extracellular lipid scavenging. We demonstrate that KRAS upregulates the LysoPC transporter MFSD2A, enabling PDAC cells to efficiently acquire extracellular unsaturated LysoPCs, maintain lipid homeostasis, sustain PI3K-AKT signaling, and promote tumor growth. Furthermore, we show that this pathway can be pharmacologically targeted using synthetic lysophospholipid analog, identifying an actionable metabolic vulnerability in PDAC.

Reduced circulating LysoPCs represent one of the most reproducible metabolic features reported in PDAC and several other malignancies. Decreased plasma LysoPC species have been observed in colorectal, breast, prostate, and pancreatic cancers, suggesting that perturbation of LysoPC homeostasis is a common feature of malignant progression [36, 37]. Consistent with our previous blinded validation study, in which LysoPC(18:0) and LysoPC(20:3) contributed to a plasma biomarker panel for early PDAC detection [12], the current cohort similarly demonstrated widespread reductions in circulating LysoPC species. Other studies further reported restoration of plasma LysoPC levels following surgical resection of PDAC, linking circulating LysoPC abundance to tumor burden [38]. While these observations have largely been interpreted in the context of biomarker discovery, accumulating evidence suggests that circulating lipids can function as metabolically important nutrients for tumors. Intravenously administered fluorescent lipid emulsions accumulated in patient-derived PDAC xenografts [39]. Restoration of circulating fatty-acid supply partially rescued PDAC growth suppressed by caloric restriction [11]. More recently, a study provided evidence that CAF can transfer blood-derived lipids to PDAC cells through membrane trogocytosis, with disruption of this process limiting tumor growth [40]. Consistent with a broader role for blood-to-tumor lipid transfer, direct movement of lipids from the circulation into tumors was demonstrated using isotope tracer approaches [41, 42]. Together, these studies support the concept that circulating and host-derived lipids are functionally important nutrient reservoirs for tumors. Our findings extend this paradigm to LysoPCs by demonstrating that circulating LysoPCs are not merely disease-associated biomarkers but also functionally important extracellular lipids that can be directly accessed and utilized by pancreatic cancer cells.

An emerging theme in PDAC biology is that extracellular nutrient acquisition is essential for adaptation to metabolic stress. Previous studies demonstrated that hypoxia and oncogenic signaling promote selective scavenging of extracellular LysoPCs [7], while cancer-associated fibroblasts (CAFs) provide unsaturated lipids that support membrane homeostasis and relieve ER stress under nutrient-limited conditions [8]. CAFs have also been identified as an important source of LysoPCs within the tumor microenvironment [8, 17]. Our findings integrate these observations into a unified model in which extracellular LysoPCs serve as transferable lipid nutrients that support PDAC growth. Notably, we observed preferential depletion and utilization of unsaturated LysoPC species, consistent with their unique value in maintaining membrane fluidity and lipid homeostasis when endogenous desaturation becomes energetically restricted. Moreover, MFSD2A expression was enriched within hypoxic tumor regions, whereas CAF-derived LysoPC production has been reported to occur predominantly in normoxic stromal compartments [8, 17]. Although direct metabolic exchange remains to be formally demonstrated, these observations support a model whereby LysoPCs produced in stromal regions become available to metabolically stressed cancer cells through MFSD2A-mediated uptake. A central finding of this study is the identification of MFSD2A as a critical mediator of LysoPC uptake in PDAC. Previous studies established MFSD2A as a sodium-dependent transporter for long-chain LysoPCs, particularly unsaturated species carrying polyunsaturated fatty acids [19, 20, 31]. However, its role in cancer has remained poorly characterized. Here, we demonstrate that MFSD2A is required for efficient LysoPC uptake, downstream LysoPC metabolism, AKT activation, and tumor cell proliferation. Importantly, MFSD2A deficiency markedly impaired the ability of extracellular LysoPC to promote growth and survival, highlighting its functional importance in PDAC metabolism. We further show that MFSD2A expression is regulated by oncogenic KRAS through MAPK signaling, providing a mechanistic explanation for earlier observations that RAS-transformed cells preferentially scavenge extracellular LysoPCs [7]. Together, these findings establish MFSD2A as a KRAS-regulated lipid transporter that expands the capacity of PDAC cells to acquire extracellular lipid nutrients under metabolically restrictive conditions.

LysoPCs are metabolically versatile molecules that function both as structural lipid intermediates and signaling mediators. Extracellular LysoPC can be reacylated into phosphatidylcholine through the Lands cycle, hydrolyzed into glycerophosphocholine and free fatty acids, or converted by autotaxin into LysoPA [35, 43, 44]. Each of these pathways has previously been implicated in PDAC progression [8, 17]. Our study reveals an additional dimension of LysoPC biology. While LysoPC-derived metabolites promoted cell growth and acute AKT activation, intact LysoPC elicited the most robust proliferative responses and uniquely sustained long-term AKT phosphorylation. The different duration in sustaining PI3K/AKT signaling suggested that LysoPA and LysoPC may activate PI3K/AKT signaling distinctly. LysoPA binds to a family of GPCRs known as LPA receptors (LPARs) [45], to induce AKT phosphorylation in minutes [46, 47]. In HeLa cells treated with LysoPA, phosphorylation of AKT can be detected as early as 5 minutes after treatment, peaked at around 30 minutes to 2 hours, and then gradually weakened [48], while in other settings, LysoPA-induced AKT phosphorylation can sustain for 24 hours [49, 50]. These findings suggest that LysoPC exerts biological activities that cannot be fully explained by its downstream metabolites alone. Although the precise mechanisms responsible for sustained AKT activation remain to be defined, our data indicate that LysoPC represents a functionally important extracellular nutrient whose activity extends beyond serving as a precursor for LysoPA or phosphatidylcholine synthesis.

Our findings also have therapeutic implications. Alkyl-lysophospholipids (ALPs) such as edelfosine are metabolically stable lysophospholipid analogs that accumulate in tumor cells and disrupt lipid-dependent survival pathways, including PI3K-AKT signaling [26, 27, 51, 52, 53, 54, 55]. Several ALPs have been evaluated in clinical studies with mixed results. Topical miltefosine produced response rates of 33.3% in a randomized trial and 22% in a phase II study of breast-cancer skin metastases [28, 56], and an overall response rate of 58% in cutaneous T-cell lymphoma [57]. However, the efficacy of ALPs in other solid tumors is very limited. In a phase II study with 81 evaluable advanced non-small cell lung cancer (NSCLC) patients, 2 had partial remissions, 68 had stable disease, and 11 had continued progression [58]. In two independent studies of advanced pancreatic cancer, the response was stable disease in one out of ten and two out of seventeen patients, respectively [29, 59]. These studies may suggest that the efficacy of ALPs is selective and determined by a yet unknown factor. ALPs are membrane-targeting agents that can be imported by cancer cells through partitioning into the outer leaflet of the plasma membrane and internalized with the help of phospholipid translocase/flippase [60]. Another study reported that ALPs enter cells in a lipid-raft-dependent manner, and disruption of raft-dependent endocytosis by knocking down of sphingomyelin synthase reduced ALPs intake and ALP-induced apoptosis [61]. The basis for this variability remains poorly understood. Our study demonstrates that MFSD2A contributes to edelfosine uptake and influences cellular sensitivity to treatment, suggesting that transporter expression may represent an important determinant of drug response. In addition, edelfosine interfered with LysoPC utilization and blocked LysoPC-induced AKT activation, indicating that therapeutic efficacy may derive, at least in part, from disruption of MFSD2A-dependent LysoPC metabolism. Although additional studies are required to define the relative contributions of MFSD2A and other mechanisms of ALP uptake, our findings raise the possibility that MFSD2A expression or LysoPC pathway activity could serve as biomarkers of target engagement and therapeutic response.

In summary, we identify a KRAS-driven metabolic adaptation in PDAC whereby oncogenic KRAS upregulates MFSD2A to enhance scavenging of extracellular LysoPCs. This pathway supports lipid homeostasis, sustains AKT signaling, and promotes tumor cell survival and growth under metabolic stress. These findings establish the KRAS-MFSD2A-LysoPC axis as a previously unrecognized component of KRAS-directed metabolic reprogramming and highlight extracellular LysoPC acquisition as a therapeutically targetable metabolic vulnerability in PDAC.

## Materials and Methods

### Cell culture

Human PDAC cell lines were obtained from American Type Culture Collection. Murine KPC and iKRAS cell lines were obtained from Dr. Maitra’s and Dr. Yao’s lab, respectively. Cells were cultured in DMEM (Cat. # MT10027CV, Corning) supplemented with 10% inactivated fetal bovine serum (FBS, Cat. #16140-071, Gibco) and maintained at 37 °C in a humidified atmosphere with 5% CO2.

### Generation of MFSD2A knockdown cells

The lentivirus-based shRNA constructs (pGIPZ) were generated by the Function Genomics Core at MD Anderson Cancer Center. The sequences of shRNA constructs (Cat. #RHS4430-200279196 and # RHS4430-200282127 for human MFSD2A; #RMM4431-200379311 and #RMM4431-200381069 for mouse MFSD2A) were obtained from Horizondiscovery (https://horizondiscovery.com/). Stable knockdown of MFSD2A in cancer cells was achieved by lentivirus infection and subsequent selection of puromycin-resistant cells and knockdown of MFSD2A was verified with qPCR.

### Lipidomic Analyses

Lipidomic analyses were performed as previously described [62]. Cells were washed 3x with ice-cold 0.9% NaCl followed by addition of 1 mL of ice-cold isopropanol:ultrapure water (3:1). Cells were scraped and transferred to a 15mL conical tube (Eppendorf), followed by briefly vortexed and centrifugation at 4°C for 10 min at 2,000 x g. Thereafter, 0.8 mL of metabolite extracts were transferred to 1.5 mL Eppendorf tubes and stored in -80°C until lipidomic analyses. For analysis of complex lipids, in a 96 well plate, 10µL (3:1 isopropanol:ultrapure water) of the cell lysates supernatant was diluted with 90µL of 1:3:2 100 mM ammonium formate, pH3: acetonitrile: 2-propanol and transferred to a 384-well microplate (Eppendorf) for analysis by LC-MS.

Untargeted lipidomic analyses were conducted on a Waters Acquity™ UPLC system coupled to a Xevo G2-XS quadrupole time-of-flight (qTOF) mass spectrometer as previously described [63, 64]. Chromatographic separation was performed using a C18 (Acquity™ UPLC HSS T3, 100 Å, 1.8 µm, 2.1×100mm, Water Corporation, Milford, U.S.A) column at 55°C. The mobile phases were (A) water, (B) Acetonitrile, (C) 2-propanol and (D) 500mM ammonium formate, pH 3. A starting elution gradient of 20% A, 30% B, 49% C and 1% D was increased linearly to 4% A,14% B, 81% C and 1 % D for 4.5 min, followed by isocratic elution at 4% A,14% B, 81%C and 1%D for 2.1 min and column equilibration with initial conditions for 1.4 min.

Mass spectrometry data was acquired using ‘sensitivity’ mode in positive and negative electrospray ionization mode within 100-2000 Da. For the electrospray acquisition, the capillary voltage was set at 1.5 kV (positive), 3.0kV (negative), sample cone voltage 30V, source temperature at 120 °C, cone gas flow 50 L/h and desolvation gas flow rate of 800 L/h with scan time of 0.5 sec in continuum mode. Leucine Enkephalin; 556.2771 Da (positive) and 554.2615 Da (negative) were used for lockspray correction and scans were performed at 0.5sec. The injection volume for each sample was 3 µL. The acquisition was carried out with instrument auto gain control to optimize instrument sensitivity over the samples acquisition time.

LC-MS and LC-MSe data were processed using Progenesis QI (Nonlinear, Waters). Annotations were determined by matching accurate mass and retention times using customized libraries created from authentic standards and/or by matching experimental tandem mass spectrometry data against the NIST MSMS or HMDB v3 theoretical fragmentations; for complex lipids retention time patterns characteristic for lipid subclasses are also considered.

To correct for injection order drift, each feature was normalized using data from repeat injections of quality control samples collected every 10 injections throughout the run sequence. Measurement data were smoothed by Locally Weighted Scatterplot Smoothing (LOESS) signal correction (QC-RLSC) as previously described.41,67 Feature values between quality control samples were interpolated by a cubic spline. Only detected features exhibiting a relative standard deviation (RSD) less than 30% in either the historical or pooled quality controls samples were considered for further statistical analysis. Metabolite values were normalized to the total ion count and reported as normalized area units.

### RNA sequencing

RNA-seq analysis was performed on KPC (shCTRL, shMFSD2A-1, shMFSD2A-2) cells (biological triplicates). RNA was extracted by RNeasy Mini Kit (Cat. #74104, Qiagen) according to the manufacturer’s protocol. RNA quality and integrity were assessed using an Agilent 2100 Bioanalyzer. All samples passed the quality control process with an RNA Integrity Number (RIN) > 7. Sequencing libraries were prepared using the Illumina TruSeq Stranded mRNA Library Prep Kit to select polyadenylated transcripts. Quality-controlled libraries were multiplexed and sequenced on an Illumina NovaSeq 6000 platform using a 2 x 150-bp paired-end configuration, aiming for a depth of 30 million reads per sample (R1+R2). Raw sequencing read quality was assessed using FastQC. High-quality reads were aligned to the mouse reference genome (GRCm39/mm39) using STAR aligner. Gene-level expression counts were quantified using featureCounts based on the GENCODE annotation. Differential gene expression analysis was performed using the DESeq2 R package. Gene Set Enrichment Analysis (GSEA) analyses were performed in R.

### Immunofluorescence

Cells were fixed with 4% paraformaldehyde and stained with p62 antibody (Cat. #39749, Cell Signaling Technology), LC3B (Cat. #83506s, Cell Signaling Technology), Alexa 488- and Alexa 647-conjugated secondary antibodies. Nuclei were stained with DAPI. Images were acquired in z-series on a spinning-disk confocal system.

### Cell viability assay

Cell viability of cancer cells was determined using the CellTiter 96 Aqueous One Solution Cell Proliferation Assay (MTS) kit (Cat. #G3580, Promega) following the manufacturer’s instructions.

### Caspase 3/7 Activity

Apoptosis activity was determined using Promega Caspase-Glo 3/7 kits (#G8210, Promega) according to manufacturer’s instructions.

### Total RNA extraction and quantitative PCR (qPCR)

Total RNA was extracted using the RNeasy Mini Kit (Cat. #74104, QIAGEN). Reverse transcription of total RNA was performed using the High-Capacity Reverse Transcriptase Kit (Cat. #4368813, Applied Biosystems). TaqMan Universal Master Mix II (Cat. #4440040, Applied Biosystems) was used for analyzing *MFSD2A* expression. Probes for MFSD2A were purchased from Thermo Fisher Scientific (Mm01192208_m1, Hs00293017_m1).

### Immunoblotting

Total proteins were extracted in RIPA buffer (Cat. #89901, Pierce) containing complete protease inhibitor cocktails (Cat. #04693116001, Roche) and phosSTOP (Cat. #04906837001, Roche). A total of 20 µg of proteins were loaded for electrophoresis in SDS-PAGE (Cat. # 1610772, Bio-Rad). Antibodies used in this study include: ACTB (Cat. #3700, Cell signaling technology), pAKT (S473) (Cat. #4060, Cell signaling technology), AKT (Cat. #9272, Cell signaling technology), p62 (Cat. #39749, Cell signaling technology), LC3B (Cat. #83506, Cell signaling technology), FASN (Cat. #3189, Cell signaling technology), SREBP1 (Cat. #PA1-337, Invitrogen), MFSD2A (Cat. #PA5-21049, Invitrogen), GAPDH (Cat. #5174, Cell signaling technology).

### In-vivo studies

Animal experiment protocols were approved by The University of Texas MD Anderson Cancer Center IRB and in accordance with the Guidelines for the Care and Use of Laboratory Animals published by the NIH (Bethesda, MD). NU/J mice (Cat. #002019, The Jackson Laboratory) were housed in specific pathogen free facilities.

For subcutaneous xenograft models of PDAC, a total of 1 x 10^5^ KPC cells were suspended in 50 µl 50% Matrigel Matrix (Corning)/Opti-MEM media and injected into 8- to 10-week-old nude mice. The dimensions of tumors were measured by a caliper, and the tumor volume was calculated by the formula V = 0.5 x length x width^2^.

### Statistical analysis

For continuous variables, statistical significance was determined by two-sided Student’s t-test unless otherwise specified. For categorical variables, statistical significance was determined by Fisher’s Exact Test for two class categorical comparisons or χ² test for trend for multiple categorical variables. Figures were generated in GraphPad Prism Version 10.0.0.

## Data availability

The data that support the findings of this study are available from the corresponding author upon reasonable request.

## Ethical approval

Animal experiment protocols were approved by The University of Texas MD Anderson Cancer Center IRB and in accordance with the Guidelines for the Care and Use of Laboratory Animals published by the NIH (Bethesda, MD).

## Acknowledgement

We sincerely thank Dr. Wantong Yao at MD Anderson Cancer Center for the gift of iKRAS cells.

---

**Supplementary Figure S1.**
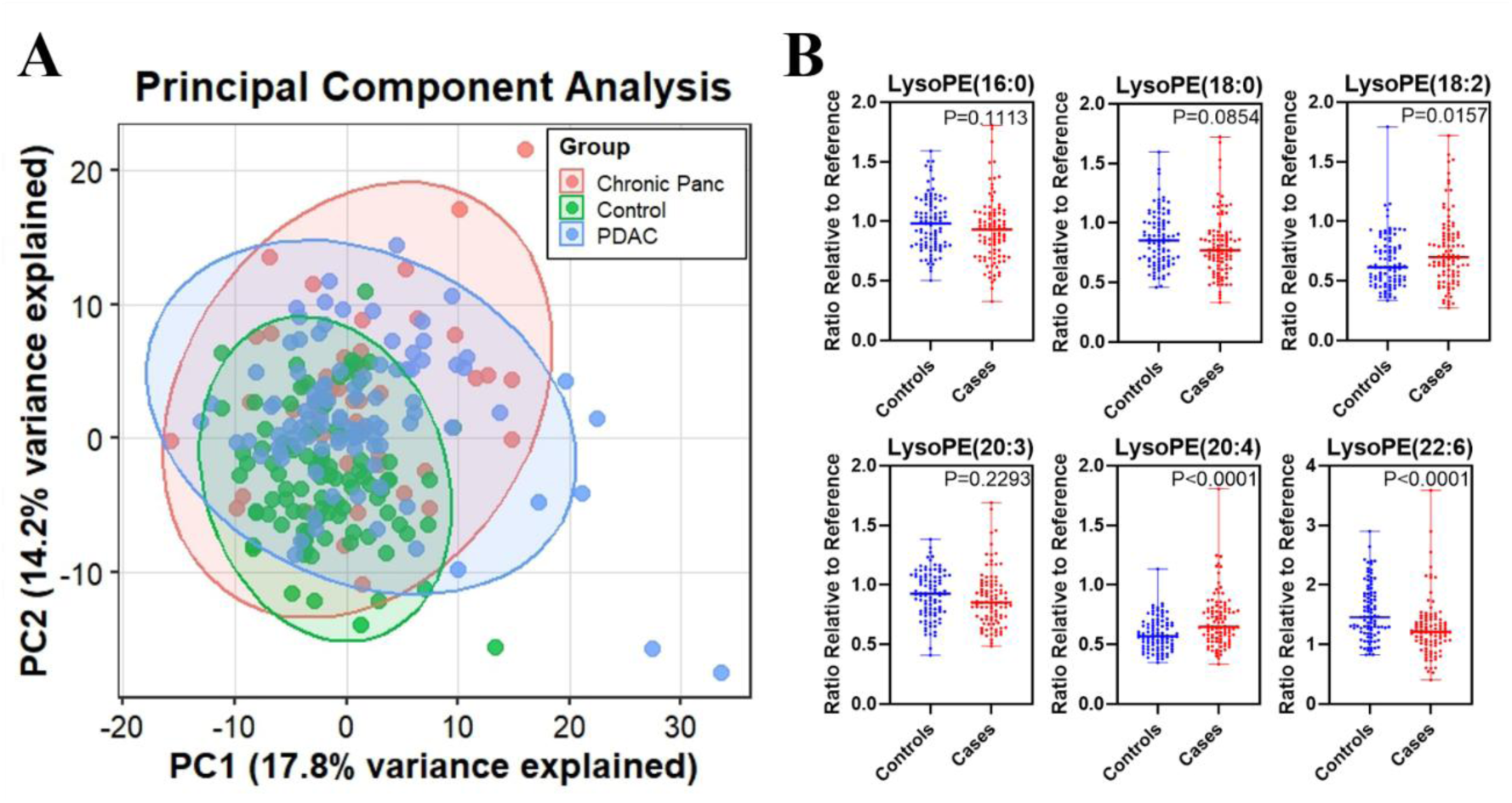
**A)** Principal component analysis (PCA) of plasma lipidomic profiles from patients with PDAC, CP, and healthy controls. Each point represents an individual plasma sample. **B**) Distribution plots illustrating relative abundance (compared to reference historical quality control) of lysophosphatidylethanolamine (LysoPE) species in plasmas of PDAC cases (n=99) and healthy controls (n=96). Statistical significance was determined by Welch’s t test and 2-sided p values are reported.

**Supplementary Figure S2.**
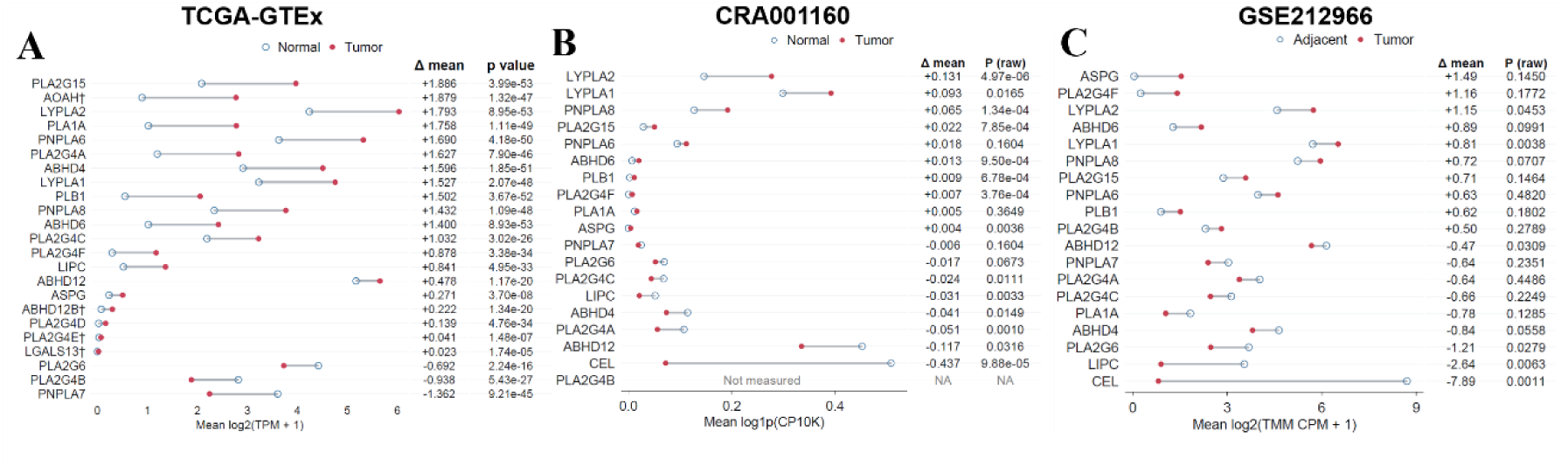
**A)** Expression levels of genes associated with LysoPC de-acylation in normal pancreas (GTEx) and PDAC (TCGA) tissues. Statistical significance was determined by two-sided Wilcoxon rank-sum test. **B-C)** Expression levels of genes associated with LysoPC de-acylation in epithelial cells of adjacent normal pancreas and PDAC tissues. Data were obtained from CRA001160 and GSE212966. Statistical significance was determined by two-sided Wilcoxon rank-sum test (CRA001160) and two-sided paired t-test (GSE212966).

**Supplementary Figure S3.**
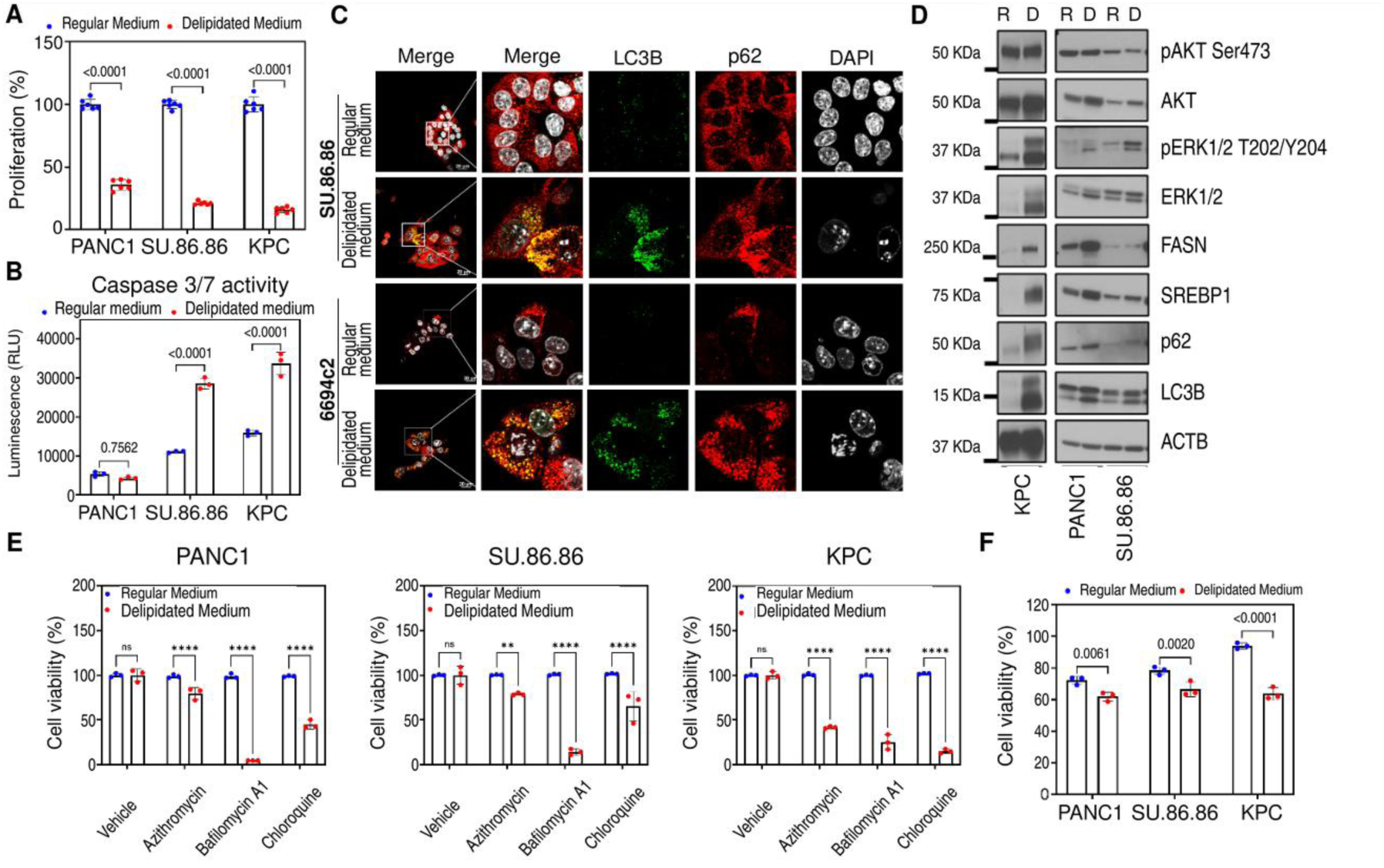
Depletion of extracellular lipids enhances *de novo* lipogenesis and autophagy. **A)** Cell viability of human PDAC and murine KPC cells cultured in regular or delipidated medium. Statistical significance was determined by Šídák’s multiple comparisons test. **B)** Caspase 3/7 activity in human PDAC and murine KPC cells cultured in regular or delipidated medium. Statistical significance was determined by Šídák’s multiple comparisons test. **C)** Confocal imaging of LC3B (Green), p62 (Red) and nucleus (white) in human PDAC and murine KPC cultured in regular or delipidated medium. **D)** Immunoblotting of phosphorylated AKT (pAKT, S473), AKT, pERK1/2 (T202/Y204), ERK1/2, FASN, SREBP1, p62, LC3B, and ACTB in human PDAC and murine KPC cultured in regular (R) or delipidated (D) medium. **E)** Cell viability of human PDAC and murine KPC cells cultured in regular or delipidated medium and treated with autophagy inhibitors. Statistical significance was determined by Fisher’s LSD. **F)** Cell viability of human PDAC and murine KPC cells cultured in regular or delipidated medium and treated with vehicle control or FASN inhibitor. Statistical significance was determined by Šídák’s multiple comparisons test.

**Supplementary Figure S4.**
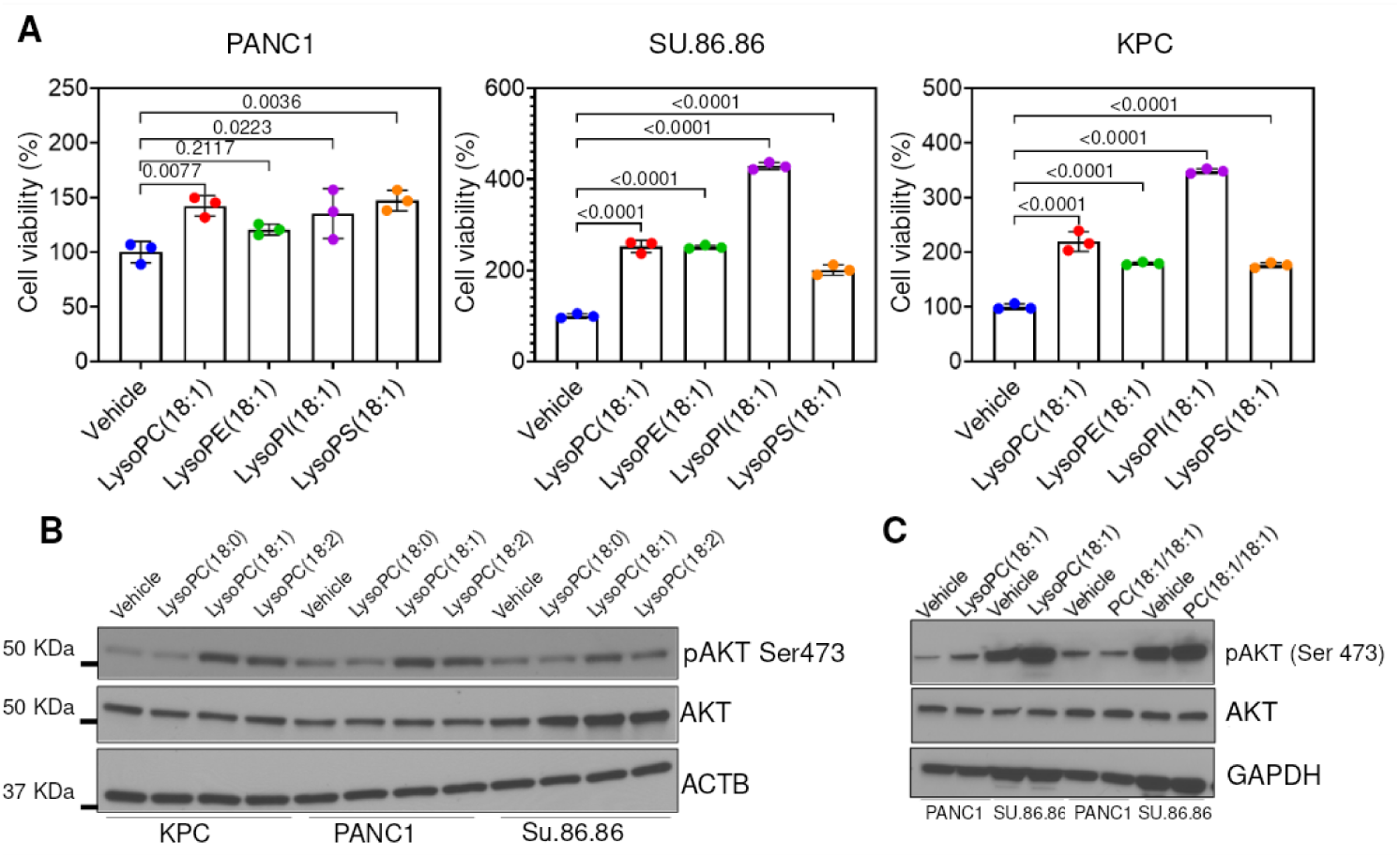
**A)** Cell viability of human PDAC and murine KPC cells treated with vehicle control, LysoPC(18:1), LysoPE(18:1), LysoPI(18:1) or LysoPS(18:1). Statistical significance was determined by Dunnett’s multiple comparisons test. **B)** Immunoblots for p-AKT (S473), AKT, and ACTB in human PDAC and murine KPC cells treated vehicle control, LysoPC(18:0), LysoPC(18:1), or LysoPC(18:2). **C)** Immunoblots for p-AKT (S473), AKT, and GAPDH in human PDAC cells treated vehicle control, LysoPC(18:1) or DOPC.

**Supplementary Figure S5.**
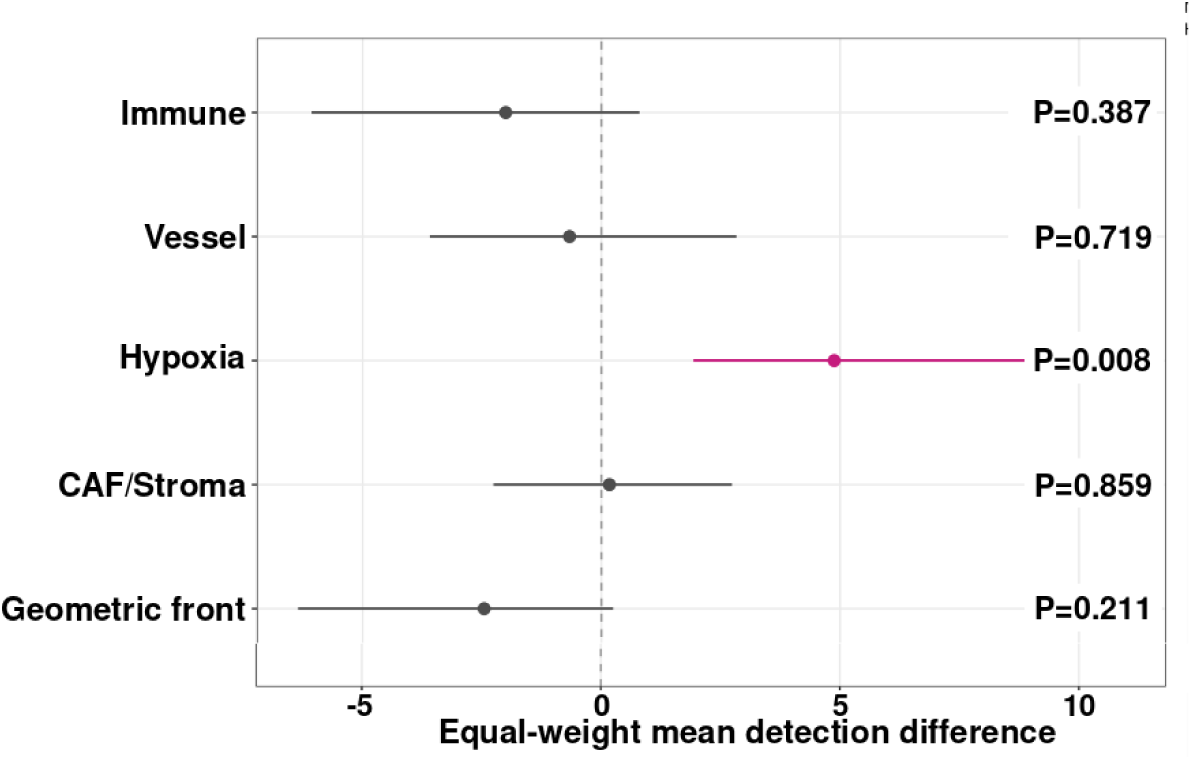
Association of *MFSD2A* expression with various annotated regions in PDAC. Data was obtained from GSE272362. Statistical significance was determined by two-sided exact section-level sign-flip test. For hypoxia, exact P=0.0078 and Benjamini-Hochberg FDR=0.039.

**Supplementary Figure S6.**
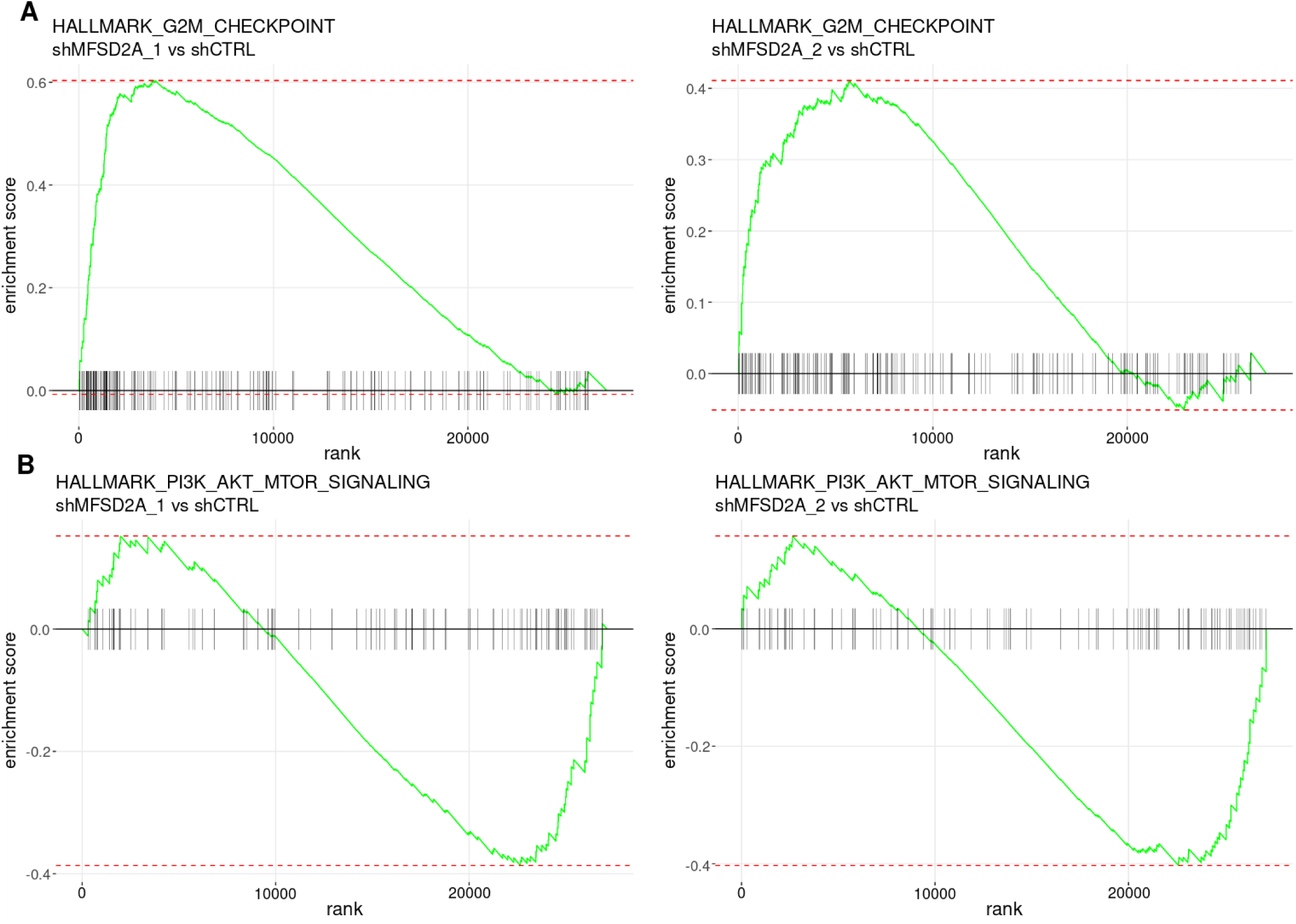
**A-B)** GSEA of Hallmark G2M checkpoint (**A**) and PI3K/AKT/mTOR (**B**) signatures in shCTRL, shMFSD2A-1, and shMFSD2A-2 cells.

